# Genomic diversity of hospital-acquired infections revealed through prospective whole genome sequencing-based surveillance

**DOI:** 10.1101/2021.10.27.466213

**Authors:** Mustapha M. Mustapha, Vatsala R. Srinivasa, Marissa P. Griffith, Shu-Ting Cho, Daniel R. Evans, Kady Waggle, Chinelo Ezeonwuka, Daniel J. Snyder, Jane W. Marsh, Lee H. Harrison, Vaughn S. Cooper, Daria Van Tyne

## Abstract

Healthcare-associated infections (HAIs) cause mortality, morbidity, and waste of healthcare resources. HAIs are also an important driver of antimicrobial resistance, which is increasing around the world. Beginning in November 2016, we instituted an initiative to detect outbreaks of HAI using prospective whole genome sequencing-based surveillance of bacterial pathogens collected from hospitalized patients. Here we describe the biodiversity of bacteria sampled from hospitalized patients at a single center, as revealed through systematic analysis of their genomes. We sequenced the genomes of 3,004 bacterial isolates from hospitalized patients collected over a 25-month period. We identified bacteria belonging to 97 distinct species, which were distributed among 14 species groups. Within these groups, isolates could be distinguished from one another by both average nucleotide identity (ANI) and principal component analysis of accessory genes (PCA-A). Genetic distances between isolates and rates of evolution varied between different species, which has implications for the selection of distance cut-offs for outbreak analysis. Antimicrobial resistance genes and the sharing of mobile genetic elements between different species were frequently observed. Overall, this study describes the population structure of pathogens circulating in a single healthcare setting, and shows how investigating microbial population dynamics can inform genomic epidemiology studies.

**Importance:** Hospitalized patients are at increased risk of becoming infected with antibiotic-resistant organisms. We used whole-genome sequencing to survey and compare over 3,000 bacterial isolates collected from hospitalized patients at a large medical center over a two-year period. We identified nearly 100 different bacterial species, suggesting that patients can be infected with a wide variety of different organisms. When we examined how genetic relatedness differed between species, we found that different species are likely evolving at different rates within our hospital. This is significant because the identification of bacterial outbreaks in the hospital currently relies on genetic similarity cut-offs, which are often applied uniformly across organisms. Finally, we found that antibiotic resistance genes and mobile genetic elements were abundant among the bacterial isolates we sampled. Overall, this study provides an in-depth view of the genomic diversity and evolution of bacteria sampled from hospitalized patients, as well as genetic similarity estimates that can inform hospital outbreak detection and prevention efforts.

## Background

Healthcare-associated infections (HAIs) affect over half a million people in the United States each year, and annual direct hospital costs for treating HAIs are estimated at over $30 billion^1–3^. A relatively small number of bacterial species account for the majority of the burden of antibiotic-resistant HAIs. Organisms belonging to the ESKAPE (*Enterococcus faecium*, *Staphylococcus aureus*, *Klebsiella pneumoniae*, *Acinetobacter baumannii*, *Pseudomonas aeruginosa* and *Enterobacter* spp.) group of pathogens are particularly problematic, due to their high burden of HAIs and frequent multidrug resistance^2,4^. In addition, while *Clostridioides difficile* is not highly antibiotic resistant, toxin-producing *C. difficile* lineages associated with significant patient morbidity and mortality have emerged in recent years, making this organism an urgent health threat^5^.

Healthcare institutions such as hospitals and long-term care facilities constitute a unique ecological niche for the proliferation and spread of antibiotic-resistant pathogens. The hospital environment has a constant flow of vulnerable populations, and widespread use of antimicrobial medications and cleaning agents provide selective pressure for the emergence and expansion of drug-resistant bacterial strains^6^. Likewise, pathogens causing HAIs possess several common biological traits that facilitate their survival and spread in healthcare environments. These traits include frequent presence and acquisition of antimicrobial resistance, asymptomatic carriage, and the ability to survive for prolonged periods on environmental surfaces such as medical equipment, or in water systems^7–9^. These factors make healthcare settings a key contributor to the increase of antibiotic-resistant bacterial infections worldwide.

Epidemiologic surveillance of HAIs requires timely and accurate ascertainment of strain type to identify patients infected with genetically related strains of the same pathogen. Surveillance using whole genome sequencing (WGS) is the gold standard for the detection of outbreaks, and has provided significant insight into the population structure of hospital-associated bacterial infections^10,11^. To improve the detection of hospital-associated transmission at our medical center, we began conducting prospective WGS surveillance of clinical bacterial isolates from hospitalized patients in November 2016, with the aim of identifying previously undetected outbreaks and characterizing pathogen transmission routes. Our approach, called Enhanced Detection of Hospital-Associated Transmission (EDS-HAT), combines prospective bacterial WGS surveillance with data mining of the electronic health record to identify outbreaks, including those that would otherwise go undetected, and their transmission routes^12–15^. In conducting this work, we have collected and sequenced the genomes of thousands of bacterial isolates. Systematic analysis of the genomes of these isolates can increase our understanding of the diversity of bacteria causing HAIs^16^.

Here we describe the genomic diversity, evolutionary rates, antimicrobial resistance gene repertoires, and mobile genetic elements carried by over 3,000 bacterial isolates sampled from patients at an academic medical center over 25 months. We uncovered a large and diverse number of species causing HAIs at our center, and showed how different population structures and evolutionary rates among these species can impact epidemiologic studies. Systematic analyses of antimicrobial resistance genes and mobile genetic elements revealed both species-specific differences as well as broader trends, and uncovered new avenues for further investigation.

## Results

### Pangenome analysis highlights the diversity of bacteria causing HAIs

The objective of this study was to use WGS to examine the genetic diversity of HAIs at a single medical center over a multi-year period, and to understand how this diversity impacts genomic epidemiology and outbreak investigations. A total of 3,004 bacterial isolates collected from 2,046 unique patients at the University of Pittsburgh Medical Center (UPMC) from November 2016 through November 2018 were sequenced and analyzed. Isolates were distributed among 14 species groups (Supplementary Tables 1 and 2, Fig. 1). The largest proportion of isolates were sampled from the respiratory tract (33.4%) followed by urinary tract (20.6%), tissue/wound (20.6%), stool (16.7%, all *C. difficile*), and blood (8.7%) (Fig. 1). The distribution of isolated species was similar between blood and tissue/wound, while the urinary tract, respiratory tract, and stool samples had different species compositions. *P. aeruginosa* was the most prevalent species isolated, with 863 isolates (28.7% of all isolates) collected from 653 unique patients. Other prevalent species included toxin-producing *C. difficile* (16.7%), methicillin-resistant *S. aureus* (MRSA, 14%) and vancomycin-resistant *E. faecalis* and *E. faecium* (VRE, 8.2%). The remaining ten species groups contained less than 200 isolates each (Supplementary Table 1). Genome sizes were highly variable, and ranged from a median length of 2.9Mb for MRSA to 7.6Mb for *Burkholderia* spp. (Fig. 2a). Pangenome collection curves constructed for genera containing multiple species showed that *Citrobacter* spp. and *Acinetobacter* spp. had the greatest pangenome diversity, perhaps due to the large number of different species sampled for these groups (Fig. 2b, Supplementary Table 2). Pangenome collection curves for individual species showed large differences in pangenome diversity between species (Fig. 2c), with MRSA and VRE *faecium* genomes having the lowest diversity, while *P. aeruginosa, C. freundii,* and *S. marcescens* had the greatest pangenome diversity of all species collected. The large and open pangenome of *P. aeruginosa* is well known^17^, however the pangenome diversity of *C. freundii* and *S. marcescens* are not well described.

**Figure 1.**
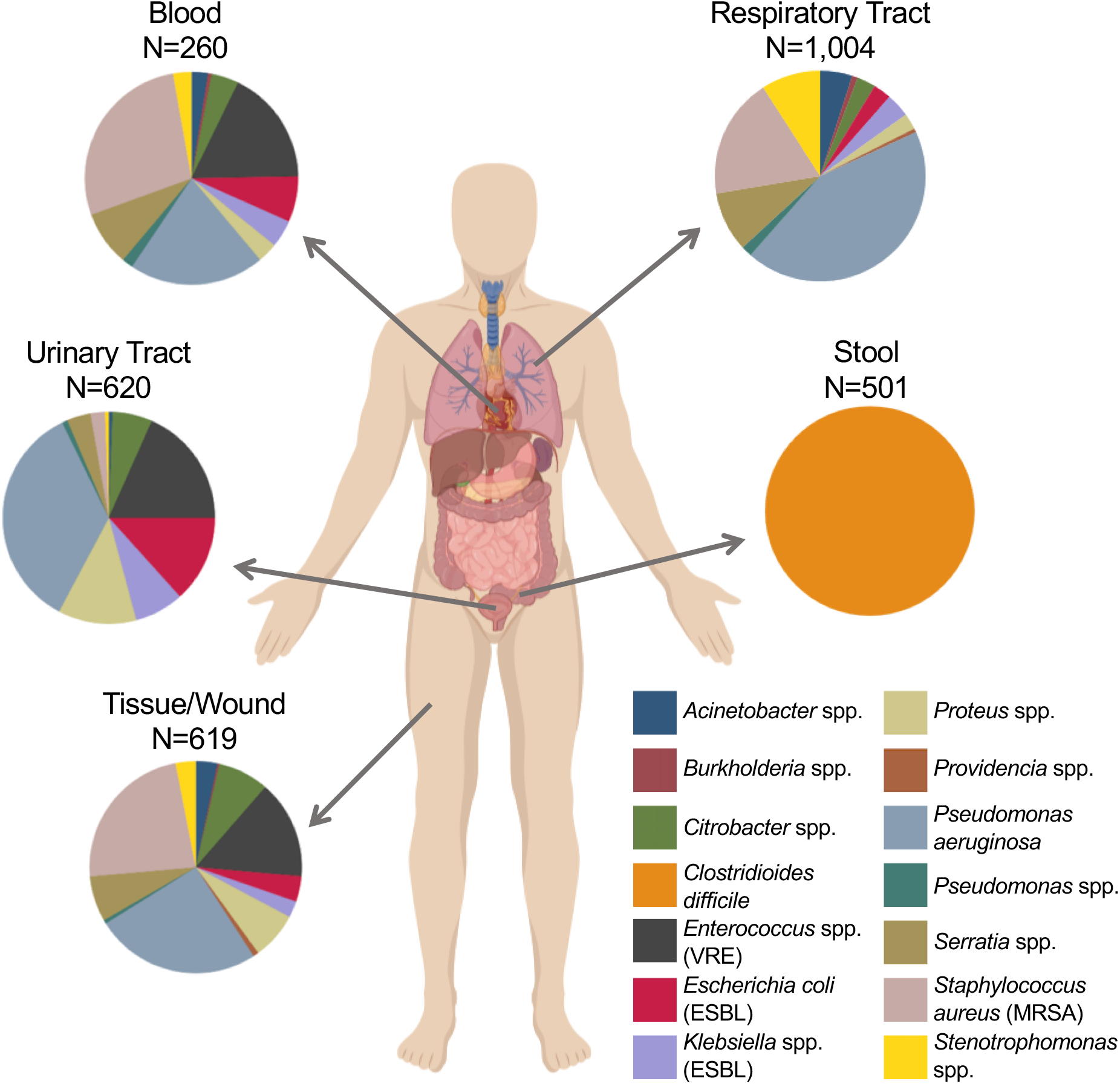
Species and body site distribution of 3,004 clinical bacterial isolates from hospitalized patients. Isolates were collected from a single hospital over 25 months as part of the Enhanced Detection System for Healthcare-Associated Transmission (EDS-HAT) project. Pie charts show the distribution of isolates belonging to 14 different species groups collected from different types of clinical specimens.

**Figure 2.**
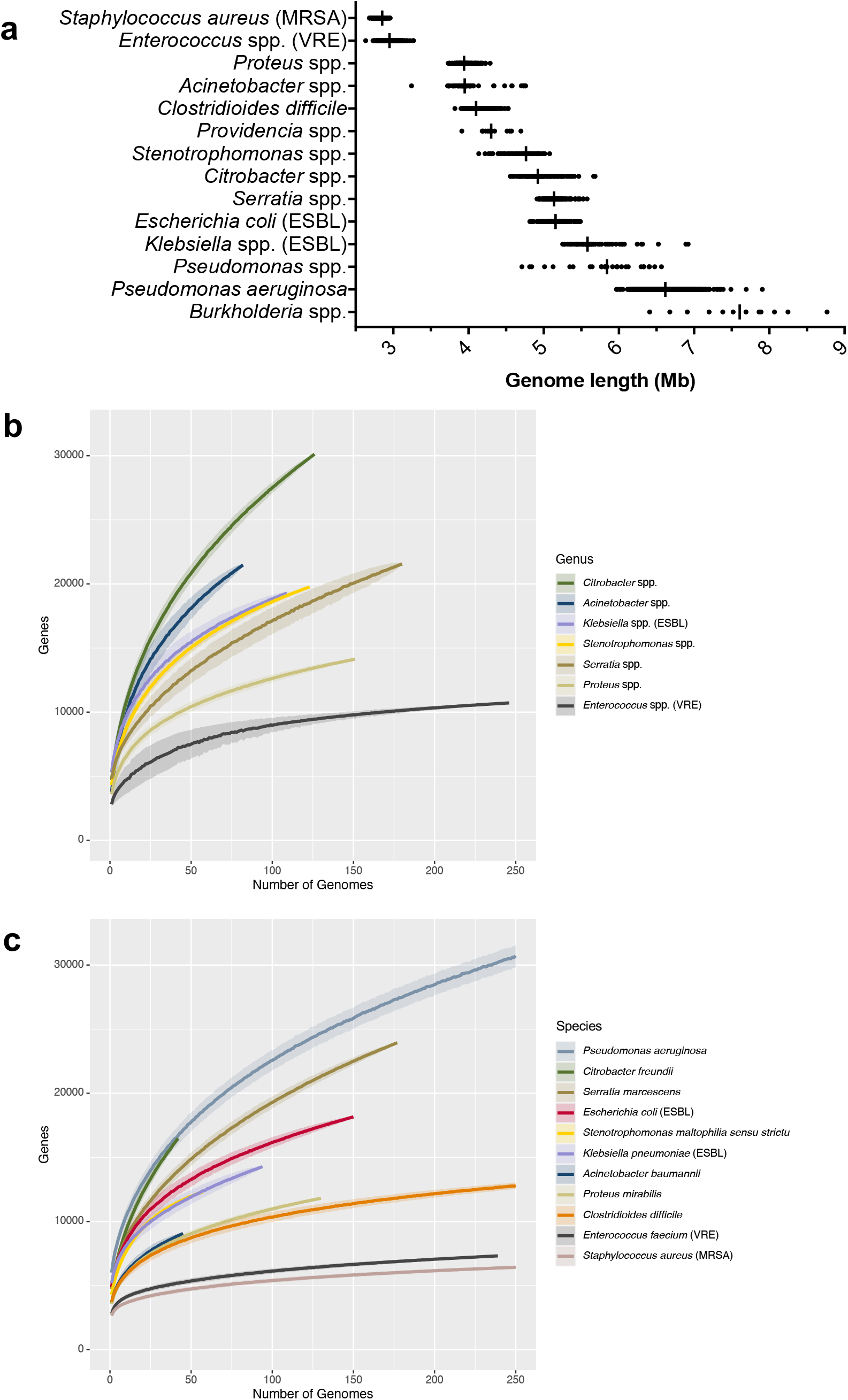
Genome length and pangenome size among sampled species. (A) Distribution of genome lengths of isolates belonging to each species group, ordered from shortest to longest median genome length. Vertical lines show median values. (B) Pangenome collection curves for up to 250 genomes from genera containing multiple species and with at least 50 genomes collected. Pangenomes were generated by Roary with an 80% protein identity cut-off. (C) Pangenome collection curves for up to 250 genomes from species with at least 40 genomes collected. Pangenomes were generated by Roary with an 95% protein identity cut-off. Curves show the mean pan-genome size and shading shows the standard deviation.

### Differences in bacterial population structures revealed by average nucleotide identity (ANI) and accessory gene content analysis

Analysis of ANI and accessory genome contents are useful methods for assigning bacterial species, as well as understanding bacterial population structures^18–20^. Because the species of each isolate collected by the EDS-HAT project was initially assigned by the clinical microbiology laboratory, we first conducted pairwise comparisons of ANI for all isolate genomes, plus additional reference genomes downloaded from the NCBI database, and used a standard 95% ANI cut-off to group genomes into the same or different species^18^. This method resulted in the identification of 97 different species among the collected isolates (Supplementary Table 2). An example of ANI-based classification of *Citrobacter* spp. is shown in Fig. 3a. As expected, several species groups were highly diverse and were composed of multiple different species, including *Acinetobacter* spp., *Burkholderia* spp., *Citrobacter* spp., *Providencia* spp., *Pseudomonas* spp., and *Stenotrophomonas* spp. (Fig. 3a, Supplementary Fig. 1). Several other species groups, such as ESBL-producing *Klebsiella* spp., *Proteus* spp. and *Serratia* spp., were composed of one dominant species (*K. pneumoniae, P. mirabilis,* and *S. marcescens*), and a small number of isolates belonging to other species (Supplementary Table 1). ANI analysis of *P. aeruginosa* identified 15 isolates (1.7% of all *P. aeruginosa* collected) that belonged to a different species and could be clearly separated from the rest of the *P. aeruginosa* population by ANI (Supplementary Fig. 2). These 15 isolates all had greater than 95% ANI with the Group 3 PA7 genome^21^, indicating that they belonged to this divergent group of *P. aeruginosa*. Overall, these findings highlight the potential discordance between species assignment based on clinical laboratory testing versus genome sequence analysis.

**Figure 3.**
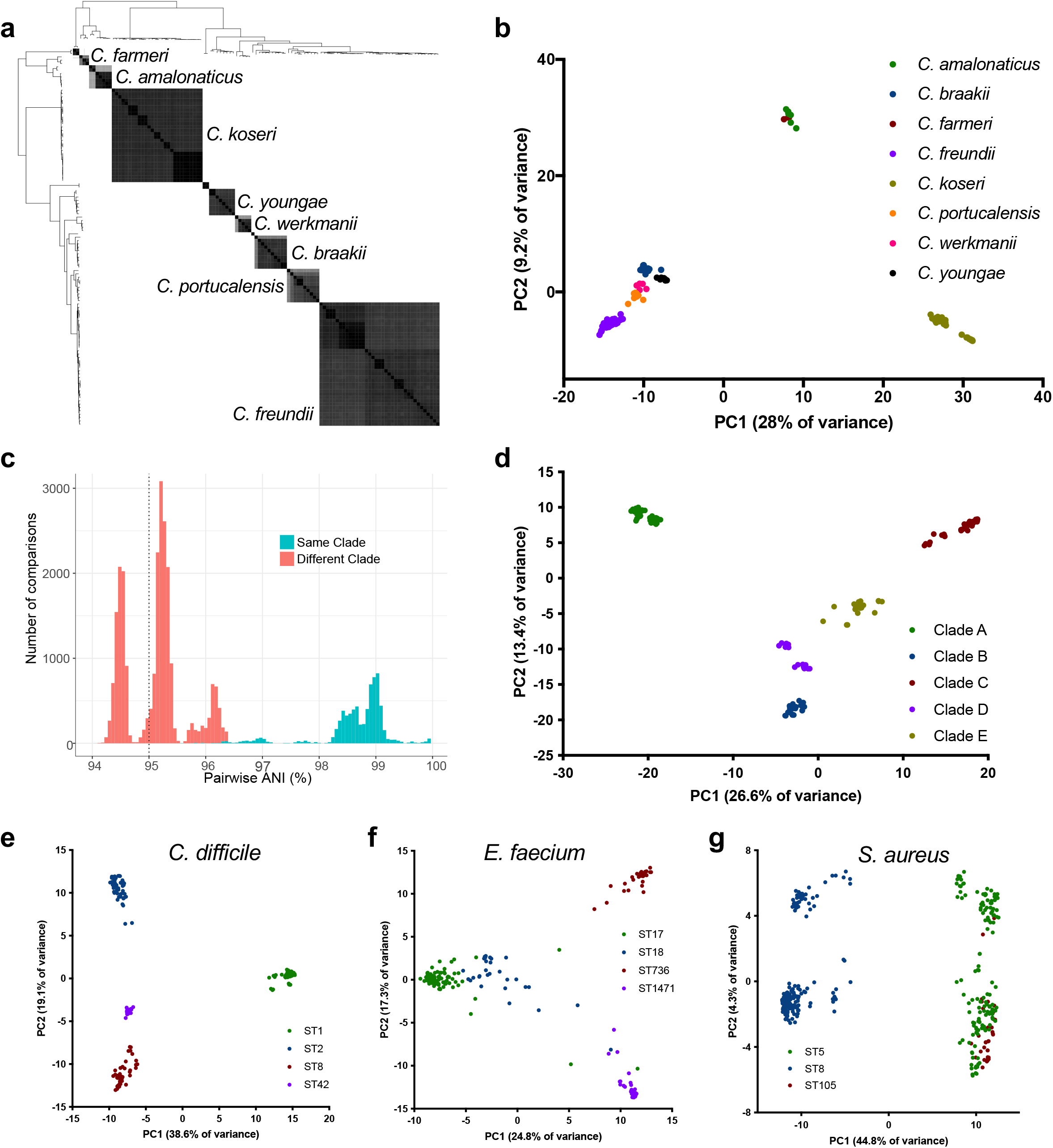
Average nucleotide identity (ANI) and principal component analysis of accessory genes (PCA-A) distinguish between and within species. (A) Phylogeny and pairwise ANI values for *Citrobacter* spp. sampled by EDS-HAT. Grey shading indicates ANI values >95%, with darker shading showing higher identity. (B) PCA-A plot for *Citrobacter* species with >2 isolates. (C) Pairwise ANI distribution of *S. marcescens* isolate genomes, showing pairwise ANI comparisons between isolates in different clades that fall below the species cut-off (95% ANI, vertical dashed line). (D) PCA-A plot for *S. marcescens* isolates, showing clear separation of five distinct clades. (E-G) PCA-A plots for dominant sequence types (STs) of *C. difficile* (E), *E. faecium* (F), and *S. aureus* (G).

While ANI measures nucleotide identity in regions that are shared between two genomes, the accessory genes, which by definition are variably present in different genomes, can also be used to identify differences between bacterial species^42,43^. We constructed principal component analysis plots based on accessory gene content (PCA-A) for species groups containing multiple species and with multiple isolates represented (Fig. 3b, Supplementary Fig. 1). The PCA-A plot for *Citrobacter* spp. isolates was largely congruent with species clustering by ANI (Fig. 3b), and the same was true for *Acinetobacter* spp. and *Stenotrophomonas* spp. as well (Supplementary Fig. 1). The *S. marcescens* isolates we collected could be clearly separated into five different clades by both ANI and PCA-A; we arbitrarily named these clades A-E (Supplementary Table 1, Supplementary Fig. 3). We observed that the pairwise ANI distribution among all *S. marcescens* isolates included comparisons of isolates in different clades that fell below the 95% ANI threshold used to distinguish species from one another (Fig. 3c, Supplementary Fig. 2). Isolates within each *S. marcescens* clade always shared greater than 95% ANI with isolates in at least one other clade, however comparisons of isolates in Clade A with isolates in either Clade C or Clade E fell below the 95% ANI threshold for same-species comparisons (Supplementary Fig. 3). PCA-A clearly separated these clades from one another (Fig. 3c), suggesting that each clade possessed a unique set of clade-specifying genes (Supplementary Table 3). These data suggest that the *S. marcescens* population we sampled may be in the process of diverging into distinct sub-species.

We also explored whether PCA-A could be used to cluster isolates belonging to different genetic lineages within a single species (Fig. 3e-g). We analyzed isolates belonging to the dominant lineages of toxin-producing *C. difficile* (Fig. 3e), VRE *faecium* (Fig. 3f), and MRSA (Fig. 3g), and found in all cases that PCA-A could generally separate isolates belonging to different STs. *C*. *difficile* isolates belonging to ST1, ST2, ST8, and ST42 were clearly separated from one another (Fig. 3e). *E. faecium* isolates belonging to ST736 were clearly separated from isolates belonging to ST17, ST18, and ST1471, which showed some overlap with one another (Fig. 3f). Finally, MRSA isolates belonging to ST8 were clearly separated from isolates belonging to ST5 and ST105, however the latter STs (which belong to the same clonal complex) were not distinguishable from one another (Fig. 3g). Analysis of gene enrichment among these different STs revealed ST-specific gene repertoires, which were largely composed of predicted mobile element genes and hypothetical proteins (Supplementary Tables 4-6). These data suggest that analysis of variable gene content may be a useful complement to SNP-based methods in epidemiologic investigations.

### Genetic diversity and evolutionary rates vary by species

The EDS-HAT project was designed to detect genetically and epidemiologically connected isolates sampled from different patients, and has successfully identified dozens of clusters containing isolates that share common exposures or transmission chains^14,15,22^. In addition, a significant number of patients in this study were repeatedly sampled. To understand how genetic diversity varied by species, we compared within-patient, within-cluster, and between-patient diversity for six different species by calculating pairwise SNP distances for all isolate pairs belonging to the same ST (Fig. 4a). In all cases, SNP differences for pairs of isolates collected from the same patient were on average lower than those for pairs of isolates collected from different patients, suggesting that patients were persistently colonized or infected with the same bacterial strain that was repeatedly sampled. Despite only comparing isolates belonging to the same ST, some same-patient comparisons for *P. aeruginosa* resulted in hundreds or thousands of SNPs, which could reflect reinfection with a different strain or the presence of hypermutator strains. Within-cluster comparisons were comparable to within-patient comparisons, demonstrating that clustered isolates were also highly genetically related to one another. We also found that there were substantial differences in median SNP distances between different species, with *C. difficile* isolates having the lowest median pairwise SNPs among isolates from the same patient (2 SNPs), and *P. aeruginosa* having the highest (15 SNPs). These data likely reflect the different genome sizes, as well as the different biology of the organisms studied here, and have broader implications for the selection of SNP cut-offs for the purposes of epidemiologic investigation.

**Figure 4.**
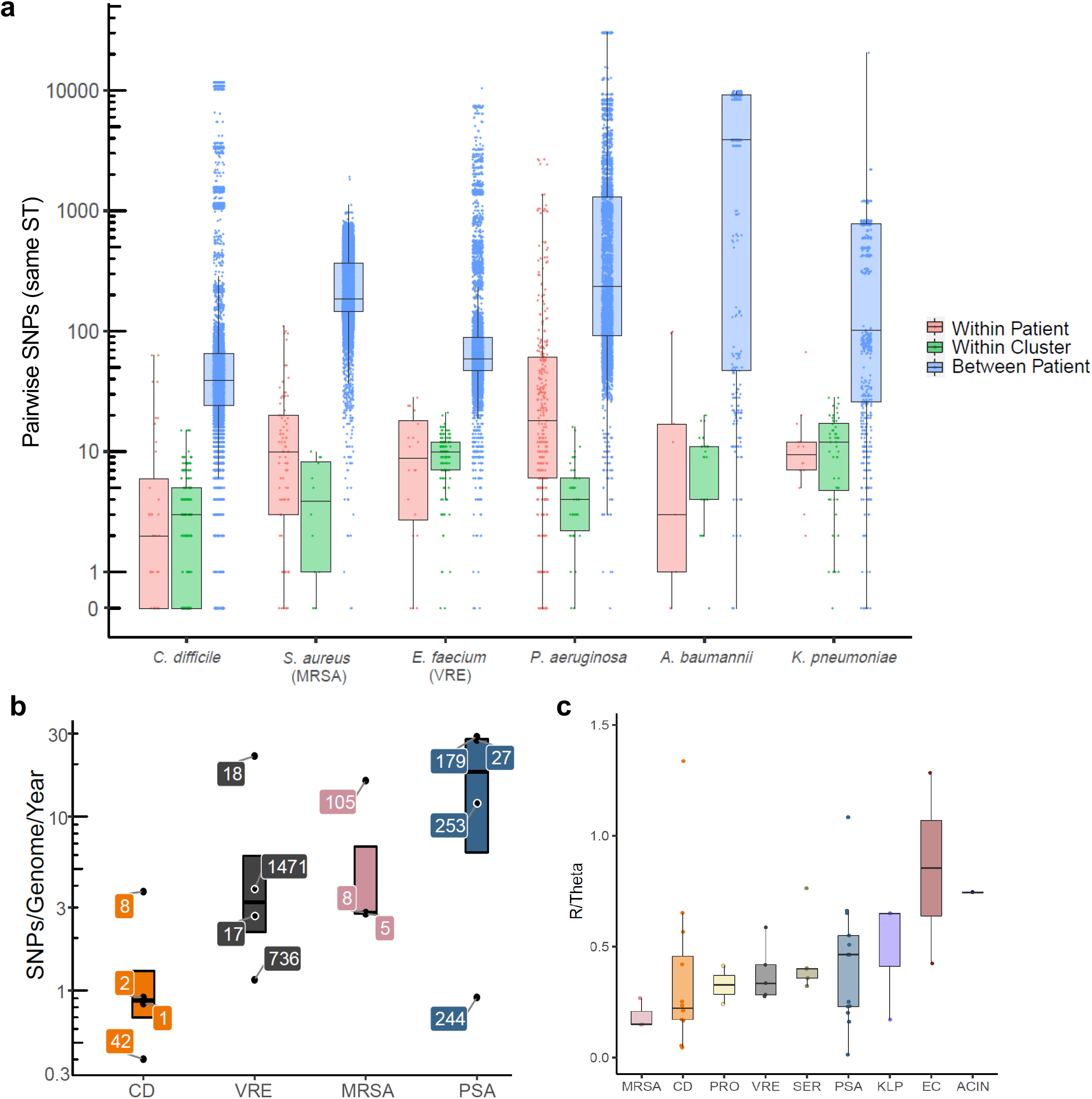
Pairwise SNP distances and genome evolution vary between species. (A) Comparison of within-patient, within-cluster, and between-patient single nucleotide polymorphisms (SNPs) for select species. Pairwise comparisons are shown for all isolate pairs belonging to the same sequence type (ST) within each species. (B) Genome evolution rates for the dominant STs within *C. difficile* (CD), vancomycin-resistant *E. faecium* (VRE), methicillin-resistant *S. aureus* (MRSA) and *P. aeruginosa* (PSA). Isolates belonging to the four largest STs (three largest for MRSA) of each species were considered, and nucleotide substitution rate (SNPs/genome/year) was calculated for each ST separately. Individual data points are labeled with the corresponding ST, and boxes show the median, 25^th^ and 75^th^ percentiles. (C) Recombination events per mutation (R/Theta) for select species. Each data point represents a distinct ST, and data are grouped by species. STs with at least 10 isolates are shown. Boxes show the median, 25^th^ and 75^th^ percentiles. PRO=*P. mirabilis*, SER=*S. marcescens*, KLP=*K. pneumoniae*, EC=*E. coli*, ACIN=*A. baumannii.*

We next compared the evolutionary rates of the *C. difficile*, VRE, MRSA, and *P. aeruginosa* populations that we sampled. We used TreeTime^23^ to estimate the nucleotide substitution rates for the most frequently observed STs for each species (Fig. 4b, Supplementary Table 7). Consistent with our observations of pairwise SNP differences (Fig. 4a), we found that *C. difficile* had the lowest evolutionary rate, VRE and MRSA had intermediate rates, and *P. aeruginosa* had the highest rate. Within each species group, however, we observed a range of nucleotide substitution rates between the different STs that were sampled. Rates overall varied nearly 100-fold among the species and STs we examined, from a minimum of 0.40 SNPs/genome/year for *C. difficile* ST42, to 28.80 SNPs/genome/year for *P. aeruginosa* ST179 (Fig. 4b, Supplementary Table 7). To understand how recombination might influence these calculations, we used ClonalFrameML^24^ to quantify the number of recombination events per point mutation (R/Theta) for each ST across all species for which at least 10 different isolates belonging to the same ST were sampled (Fig. 4c). MRSA genomes were found to have the lowest rates of recombination, while *K. pneumoniae*, *E. coli*, and *A. baumannii* appeared to have the highest rates. These data show that rates of nucleotide substitution and recombination are variable across STs as well as across species; this variability should be considered when assessing genomic similarity between isolates during epidemiologic investigations.

### Systematic analysis of antimicrobial resistance (AMR) genes uncovers broad and species-specific trends

AMR threatens the effective treatment and prevention of bacterial infections. To understand the diversity and distribution of AMR genes among the 3,004 isolates we sampled, we identified resistance genes within each genome by querying the ResFinder database with BLASTn^25^ (Supplementary Figure 4, Supplementary Table 8). The total number of AMR genes identified per genome ranged from 0-19, with an average of 4.6 AMR genes per genome. The species groups carrying the most AMR genes were *Klebsiella* spp. (average 13.1 AMR genes per genome), *E. coli* (7.7 AMR genes per genome), and VRE (average 7.4 AMR genes per genome) (Supplementary Table 8). We also classified each AMR gene by drug class, and examined the distribution of AMR genes found in more than one species group (Fig. 5a). Several genes encoding aminoglycoside and sulfonamide resistance were observed in the majority of different species groups, suggesting that AMR genes for these antibiotic classes are relatively widespread among bacterial pathogens within our hospital. The Gram-positive species we collected (*C. difficile*, VRE, and MRSA) carried different AMR genes compared to the sampled Gram-negative species, and all Gram-positive species were found to carry the aminoglycoside resistance genes *aac(6’)-aph(2’)* and *aph(3’)-III* and the tetracycline resistance gene *tet(M)*, albeit at varying frequencies (Fig. 5a).

**Figure 5.**
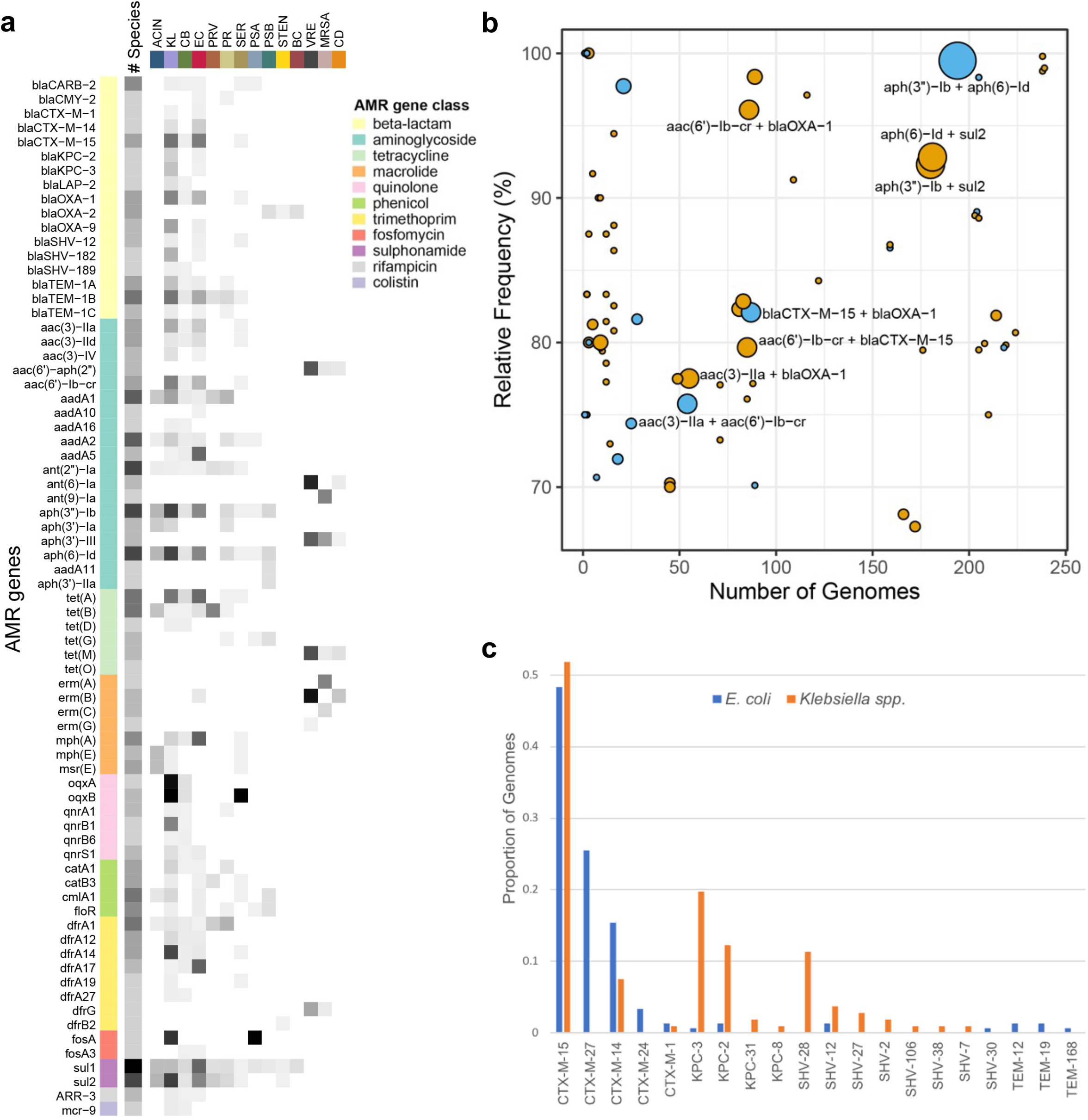
Antimicrobial resistance gene abundance and diversity. (A) Prevalence of resistance genes found in more than one species group. Genes are grouped by antibiotic class, and grey shading shows the prevalence of each gene within and across each group. Darker shading indicates higher prevalence. ACIN=*Acinetobacter* spp.; KL=*Klebsiella* spp.; CB=*Citrobacter* spp.; EC=*E. coli*; PRV=*Providencia* spp.; PR=*Proteus* spp.; SER=*Serratia* spp.; PSA*=P. aeruginosa*; PSB*=Pseudomonas* spp.; STEN*=Stenotrophomonas* spp.; BC*=Burkholderia* spp.; VRE=vancomycin-resistant *Enterococcus* spp.; MRSA=methicillin-resistant *S. aureus*; CD=*C. difficile*. (B) Resistance gene co-occurrence. Relative frequency versus number of genomes is plotted for pairs of resistance genes that co-occur at ≥50% relative frequency. Blue dots indicate AMR genes in the same drug class, while orange dots indicate genes in different classes. The size of each dot corresponds to the number of different species groups found to carry each pair. AMR gene pairs found in ≥4 different species groups are labeled. (C) Distribution of extended-spectrum beta-lactamase (ESBL) and carbapenemase enzymes among *E. coli* and *Klebsiella* spp. isolates.

We next examined the co-occurrence of pairs of AMR genes across different species groups (Fig. 5b). We found that the aminoglycoside resistance genes *aph(3”)-Ib* and *aph(6)-Id* were almost always found together, and co-occurred in eight different species groups (all Gram-negative species groups except for *Burkholderia* spp., *Providencia* spp., and *Stenotrophomonas* spp.). Both of these genes also frequently co-occurred with the sulfonamide resistance gene *sul2* (Fig. 5b). A separate aminoglycoside resistance gene, *aac(6’)-Ib-cr*, was found to frequently co-occur with the narrow-spectrum beta-lactamase *bla*_OXA-1_ as well as with the extended-spectrum beta-lactamase (ESBL) *bla*_CTX-M-15_. Finally, we examined the distribution of ESBL and carbapenemase enzymes among the ESBL-producing *E. coli* and *Klebsiella* spp. isolates that we sampled (Fig. 5c). The most frequently observed ESBL enzyme was CTX-M-15, which was found in roughly half of all *E. coli* and *Klebsiella* spp. genomes (Fig. 5c). The other half of isolates within each species group carried largely different enzymes from one another, with most *E. coli* isolates carrying other CTX-M-type and a small number of TEM-type ESBLs, while *Klebsiella* spp. isolates carried CTX-M-14 and SHV-type ESBLs. The carbapenemases KPC-2, KPC-3, KPC-8, and KPC-31 were found almost entirely among *Klebsiella* spp. genomes (Fig. 5c). These data highlight the abundant diversity of AMR genes carried by the bacteria in our hospital, and can be useful for developing tailored treatment and prevention approaches for different bacterial pathogens.

### Mobile genetic element (MGE) distribution and cargo

MGEs are frequently found within the genomes of bacteria residing in the hospital environment, and they often encode useful functions like AMR and virulence factors^26^. To assess the presence of MGEs in our dataset in a systematic and unbiased manner, we used a previously developed approach to identify nucleotide sequences with high homology (>99.9% identity over at least 10Kb) that were present in genomes of different genomospecies^27^ (Fig. 6a). This approach resulted in the identification of 186 clusters of shared sequences, which were present in 805 (26.8%) of the genomes in our dataset (Fig. 6b). While each of the 14 different species groups we sampled contained at least one genome encoding a shared sequence, species groups that were particularly enriched for shared sequences included *Klebsiella* spp., *P. aeruginosa*, and *Stenotrophomonas* spp. (Fig. 6b). We next used comparisons with available MGE databases and manual curation to assign an MGE type to each of the 186 clustered sequences based on sequence homology to previously described MGEs (Fig. 6c). We identified similar numbers of sequences that resembled insertion sequences (ISs) or transposons and that resembled prophages or integrative conjugative elements (ICEs). Slightly more sequences showed homology to plasmid sequences, and a large number of sequences resembled multiple MGE types (Fig. 6c). Importantly, 53 (28.5%) shared sequence clusters could not be assigned to an MGE type. Some of these sequences are likely fragments of larger MGEs that lacked genetic features that would enable their classification. Alternately, some of these may constitute novel MGEs.

**Figure 6.**
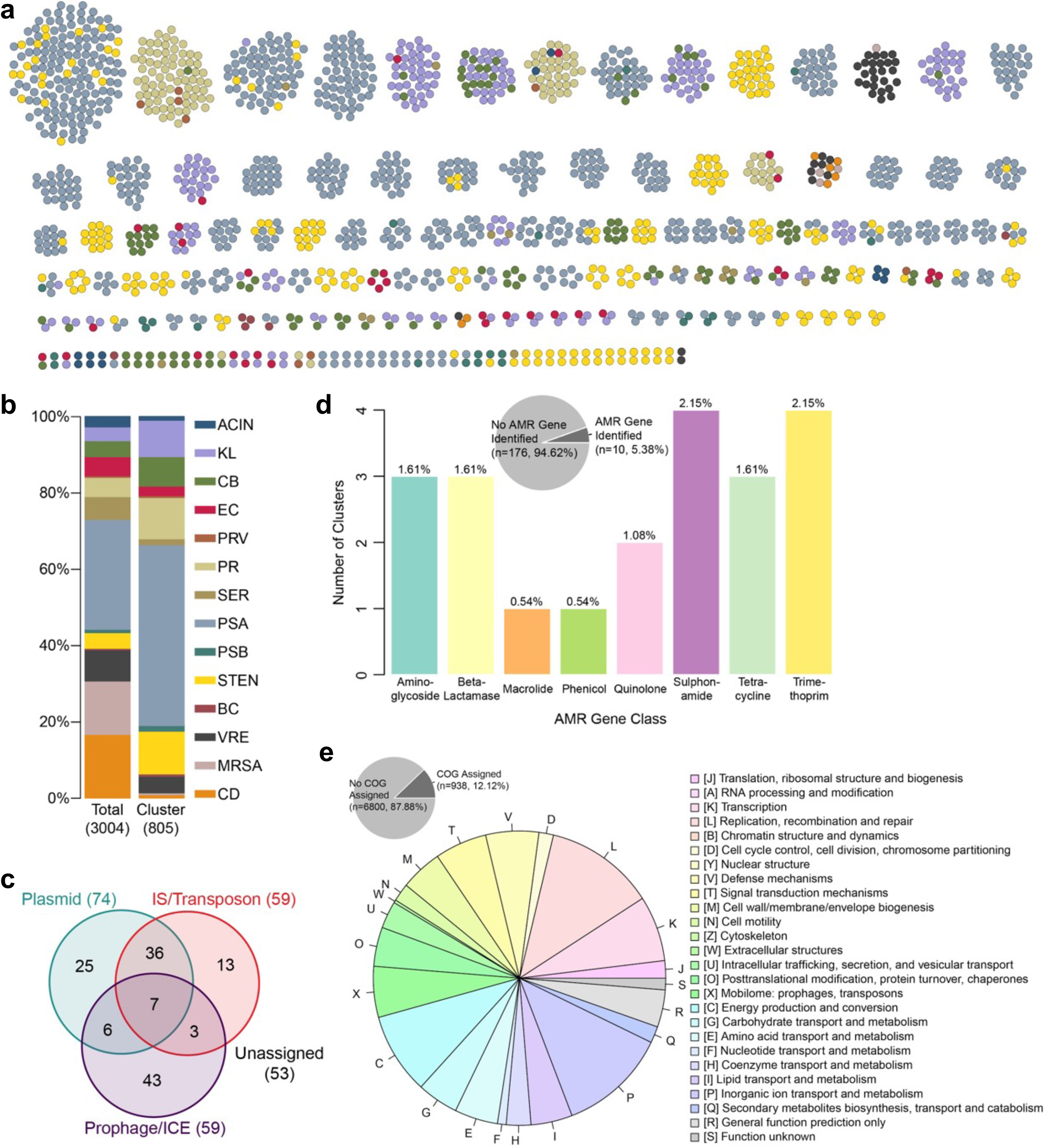
Mobile genetic element (MGE) distribution and cargo. (A) Clusters of putative MGEs identified in 3,004 study isolate genomes. Nodes within each cluster correspond to bacterial isolates, and are color coded by species group (color key provided in panel B). (B) Distribution of isolates in the entire dataset (left) versus isolates encoding one or more putative MGEs (right). (C) Distribution of putative MGEs resembling plasmid, IS/transposon, or prophage/ICE sequences, determined by nucleotide sequence comparisons and manual curation. (D) Distribution of antimicrobial resistance (AMR) genes detected among 186 putative MGEs. (E) Distribution of clusters of orthologous groups of proteins (COG) categories of MGE genes with COG categories assigned.

To understand more about the cargo encoded by the putative MGEs we identified, we first assessed the distribution of AMR genes among the 186 shared sequence clusters we studied (Fig. 6d and Supplementary Table 9). Only 10/186 shared sequence clusters (5.4%) carried AMR genes, however these clusters were found among 116/805 isolates (14.4%). The most frequently observed AMR gene classes (which were each only present in four shared sequence clusters) were sulfonamide and trimethoprim resistance, while aminoglycoside resistance genes, tetracycline resistance genes, and beta-lactamases were each found in three shared sequence clusters (Fig. 6d). We next examined the distribution of clusters of orthologous groups of proteins (COG) categories among all genes present in all shared sequence clusters in our dataset. A total of 938 genes (12.1% of all shared sequence cluster genes) had COG categories assigned, and among these genes the two COG categories observed most frequently were genes involved in replication, recombination and repair, and genes involved in inorganic ion transport and metabolism (Fig. 6e). These data suggest that prominent cargo among the shared sequences we identified included genes for MGE maintenance and transmission, as well as genes required for the utilization of and resistance to heavy metals, which pathogens frequently encounter in the hospital environment^28^.

## Discussion

HAIs place a large burden on healthcare systems by increasing patient morbidity, mortality, and the cost of medical care. The broader aim of the EDS-HAT project is to improve the detection of bacterial outbreaks in hospitals, and the project has been successful in this regard^14,15,22^. The EDS-HAT project has also provided a large dataset of microbial genomes sampled from thousands of patients within a single medical center over time. Here we highlight the genetic diversity among bacterial pathogens causing HAIs; understanding this diversity can better inform genomic epidemiology and outbreak investigations. As bacterial WGS becomes increasingly routine in healthcare settings, this study also provides a baseline for future comparisons, both at our center and elsewhere.

Using comparative genomics methods, we revealed the vast diversity among bacterial pathogens within our hospital. We identified bacteria belonging to 97 different species, which spanned 14 different species groups. We also identified 23 species which have not been previously described, including potentially novel species of *Acinetobacter, Citrobacter, Proteus, Providencia, Pseudomonas, Serratia* and *Stenotrophomonas*. A total of 41 isolates (1.4% of sampled isolates) belonged to these novel species, which was a lower proportion than that observed in a prior study of HAIs among ICU patients conducted in 2015^16^. This could be due to additional species having been described in recent years, as well as different inclusion criteria and study populations between the prior study and our own. Further investigation into these new species can aid in the clinical diagnosis of bacteria causing infections.

Our finding that both ANI and PCA-A are effective at distinguishing between different groups at both the species and sub-species levels is consistent with prior studies^29,30^. The 15 *P. aeruginosa* isolates we identified as having 93-94% ANI with the remaining *P. aeruginosa* population is also consistent with prior reports of the *P. aeruginosa* population^31^. Conversely, *S. marcescens* is known to have a population structure comprised of multiple clades^32,33^, however we found that pairwise comparisons between some of these clades had less than 95% ANI, suggesting a large degree of divergence and possible ongoing sub-speciation. We were also able to use accessory gene content differences to distinguish between the dominant genetic lineages of *C. difficile*, VRE *faecium*, and MRSA. Further investigation of these accessory genes would likely enhance our understanding of how different genetic lineages are able to co-exist in the same hospital, and could provide useful biomarkers for tracking lineages of interest.

Comparing within-patient versus between-patient genetic diversity can provide important guidance in defining SNP cut-offs for outbreak investigations. We found that the number of SNPs among genomes isolated from the same patient at different time points varied by species, with within-patient SNPs being lowest for *C. difficile*, moderate for MRSA and VRE, and greatest for *P. aeruginosa*. Differences between species likely reflect both genome size as well as the biology of these organisms; for example, *C. difficile* can spend long periods of time in a non-replicative spore state, while *P. aeruginosa* genomes are more than double the size of MRSA and VRE genomes. The SNP distances among same-patient isolates we observed are comparable to those used in outbreak investigations in our setting and elsewhere^14,34,35^. These data demonstrate that same-patient genome pairs can be used to empirically determine genetic similarity thresholds for genomic epidemiology purposes. Evolutionary rates assessed for the four most common species in our hospital were also consistent with previous studies^36,37^. The large variability in evolutionary rates between different species, however, further suggests that different SNP cut-offs should be considered for different bacterial species for the purposes of hospital outbreak investigations.

This study establishes the diversity of antimicrobial resistance genes among pathogenic bacteria circulating at our hospital, and provides a point of comparison with other studies of antibiotic resistance spread in the hospital environment^22,27,38,39^. We found that aminoglycoside and sulfonamide resistance genes were highly abundant, and were found in the majority of species that we sampled. Although the presence of aminoglycoside resistance is well documented among both Gram-positive and Gram-negative bacteria—and more specifically among the ESKAPE pathogens—less attention has been focused on sulfonamide resistance^40–42^. The co-occurrence of *aph(3”)-Ib*, *aph(6)-Id,* and *sul2* has been previously observed in a variety of different genetic contexts, including in plasmids, integrative conjugative elements, and chromosomal genomic islands^41,43^. Additionally, we found that the ESBL enzyme *bla*_CTX-M-15_ was widely distributed among both *E. coli* and *Klebsiella* spp. isolates, which is consistent with prior reports^44^. Among the other ESBL-producing *E. coli* and *Klebsiella* spp. isolates collected, ESBL enzymes were largely restricted to one species group or the other. Finally, while we did not explicitly collect carbapenemase-producing organisms during this study period, a subset of the ESBL-producing *E. coli* and *Klebsiella* spp. isolates collected also carried carbapenemase enzymes. Co-occurrence of ESBL enzymes and carbapenemases was more frequent among *Klebsiella* spp., especially ST258 *K. pneumoniae*^22^.

This study also offers an overview of highly similar sequences (which we suspect largely belong to MGEs) shared among the genomes of distantly related bacteria sampled from patients residing in the same hospital environment. We found that *Enterobacteriaceae* such as *Klebsiella* spp. and *Citrobacter* spp., as well as *P. aeruginosa* and *Stenotrophomonas* spp., were overrepresented among shared sequence clusters compared to their overall distribution in the dataset. Most of the shared sequences identified in *Enterobacteriaceae* genomes resembled sequences carried on plasmids, consistent with the frequent plasmid exchange known to happen among species in this family^45^. On the other hand, many of the shared sequences identified among *P. aeruginosa* and *Stenotrophomonas* spp. resembled prophages and integrated conjugative elements, suggesting that these organisms may rely on different MGEs to exchange genetic material. Somewhat surprisingly, our analysis identified fewer shared sequences carrying AMR genes compared to a prior study we conducted within the same hospital^27^. This may be due to our use of a longer sequence length cut-off for shared sequence identification in this study, as AMR genes are known to be carried on smaller MGE units that can rapidly shuffle, interchange, and mutate^46^. Finally, we found it notable that genes encoding metal transport and resistance were frequently observed within the shared sequences we identified. Inorganic ions are required for catalysis of many bacterial enzymes^47^, and heavy metals such as silver, copper, and mercury have long been used as disinfectants in hospitals^48^. Further study of MGEs encoding metal-interacting genes will be a focus of our future work.

This study had several limitations. The organisms we collected were pre-specified, and certain groups, such as *Enterobacter* spp. or carbapenemase-producing organisms without a noted ESBL phenotype, were not collected. Furthermore, our definition of “hospital-acquired infections” was quite broad; some of the collected isolates likely represent commensal organisms or pathogen colonization, rather than true infection. We also cannot say for sure whether the sampled bacteria were acquired from the healthcare setting or not, as we only considered bacterial isolates from clinical specimens and did not include environmental sampling. Additionally, our 25-month collection window was quite short, thus we were unable to draw conclusions regarding trends over time. Finally, the inclusion of both broad species groups as well as more defined sets of specific pathogens made it difficult to conduct systematic analyses or draw broader conclusions across the entire dataset. Nonetheless, the large number of isolates collected offers a high-resolution view of the genomic diversity and evolution of important bacterial pathogens found within our hospital. Our future work will include following these bacterial populations over time, and comparing our results with similar studies conducted in other settings.

In assessing the genomes of major infection-associated bacterial species isolated from patients at our hospital, we have provided a longitudinal survey of the genomic diversity of bacterial HAIs at a single clinical center. Our findings demonstrate that studying population dynamics and evolution of these pathogens can inform genomics-based outbreak investigations. In addition to forming a basis for future comparisons, this study also provides a deeper understanding of the breadth of different species that cause HAIs, and demonstrates the utility of systematic genome sequencing and comparative genomics analysis of clinical bacterial isolates from hospitalized patients.

## Methods

### Isolate collection

Bacterial isolates were collected from the University of Pittsburgh Medical Center (UPMC) Presbyterian Hospital, an adult tertiary care hospital with over 750 beds, 150 critical care unit beds, more than 32,000 yearly inpatient admissions, and over 400 solid organ transplants per year. Isolates were collected from November 2016 through November 2018 from admitted patients as part of a prospective genomic epidemiology surveillance project called Enhanced Detection System for Healthcare-Associated Transmission (EDS-HAT). Inclusion criteria were hospital admission greater than two days before the culture date, and/or a recent inpatient or outpatient UPMC hospital encounter in the 30 days before the culture date. A total of 3,004 isolates were included in this study (Table S1). The EDS-HAT project collected all organisms meeting the above inclusion criteria and belonging to the following genera: *Acinetobacter* spp.*, Burkholderia* spp., *Citrobacter* spp., *Proteus* spp., *Providencia* spp., *Pseudomonas* spp. *Serratia* spp., and *Stenotrophomonas* spp. Isolate collection was limited to only toxin-producing strains of *Clostridioides difficile*, vancomycin-resistant *Enterococcus* spp. (VRE), extended-spectrum beta-lactamase (ESBL)-producing *Escherichia coli* and *Klebsiella* spp., and methicillin-resistant *Staphylococcus aureus* (MRSA). This study was approved by the University of Pittsburgh Institutional Review Board and was classified as being exempt from patient-informed consent.

### Whole genome sequencing and genome assembly

Genomic DNA was extracted from pure overnight cultures of single bacterial colonies using a Qiagen DNeasy Tissue Kit according to the manufacturer’s instructions (Qiagen, Germantown, MD). Illumina library construction and sequencing were conducted using an Illumina Nextera DNA Sample Prep Kit with 150bp paired-end reads, and libraries were sequenced on the NextSeq 550 sequencing platform (Illumina, San Diego, CA). Selected isolates were re-sequenced with long-read technology on a MinION device (Oxford Nanopore Technologies, Oxford, United Kingdom). Long-read sequencing libraries were prepared and multiplexed using a rapid multiplex barcoding kit (catalog SQK-RBK004) and were sequenced on R9.4.1 flow cells. Base-calling on raw reads was performed using Albacore v2.3.3 or Guppy v2.3.1 (Oxford Nanopore Technologies, Oxford, UK).

Genome sequence analyses were performed on a BioLinux v8 server^49^ using publicly available genomic analysis tools wrapped together into a high-throughput genome analysis pipeline. Briefly, Illumina sequencing data were processed with Trim Galore v0.6.1 (https://www.bioinformatics.babraham.ac.uk/projects/trim_galore/) to remove sequencing adaptors, low-quality bases, and poor-quality reads. Kraken v1^50^ taxonomic sequence classification of raw reads was used to confirm species designation, and to rule out contamination. Illumina reads were assembled with SPAdes v3.11^51^. Long-read sequence data generated for other studies^22,27,39^ were combined with Illumina data for the same isolate, and hybrid assembly was conducted using unicycler v0.4.7 or v0.4.8-beta^52^. Assembled genomes were annotated using Prokka v1.14 and assembly quality was verified using QUAST^53^. Genomes were included in the study if they had at least 35-fold Illumina read coverage, had assemblies with ≤ 350 contigs, and had total genome lengths ± 25% of the median of all isolates within each species group. Antimicrobial resistance and toxin genes were confirmed using BLASTn in line with EDS-HAT study phenotypic inclusion criteria. Specifically, all *S. aureus* genomes were confirmed to encode the *mecA* gene, all *E. faecalis* and *E. faecium* genomes were confirmed to encode a VanA or VanB operon, all *E. coli* and *Klebsiella* spp. genomes were confirmed to encode an identifiable extended-spectrum beta-lactamase (ESBL) enzyme, and all *C. difficile* genomes were confirmed to encode either toxin A and/or toxin B genes.

### Classification of genomospecies and lineages

Within each species group, genome assemblies from this study and reference genome assemblies downloaded from the NCBI RefSeq database underwent pairwise average nucleotide identity (ANI) analysis using FastANI v1.3^18^. Genomes with ANI values >95% then underwent single-linkage hierarchical clustering using the hclust function from the R package stats v3.6. Each ANI cluster was manually assessed and assigned to a species based on the predominant nomenclature of genomes of type/reference strains within each cluster. Clusters that did not contain reference genomes, or where reference genomes were only named at the genus level, were named “genomospecies.” Sequential numbers were appended to each uncharacterized genomospecies within a species group. Species identified using ANI and having greater than 100 isolates were further sub-divided into clades and lineages based on multi-locus sequence typing (ST), or phylogenetic analysis. STs were determined from assembled contigs using mlst v2 (https://github.com/tseemann/mlst). Species without a defined ST scheme (*P. mirabilis* and *S. marcescens*) were classified into clades or lineages by grouping isolates that shared <1000 core genome single nucleotide polymorphism (SNP) differences into the same lineage, with SNPs identified using snippy (https://github.com/tseemann/snippy). *Stenotrophomonas* genomospecies were named according to Gröschel et al.^54^.

### Gene content and pangenome analyses

Gene content matrices were obtained for all species groups with more than 50 isolates using the pangenome analysis program roary v3.11^55^. Roary was run using a protein identity cut-off of 80% for genera containing multiple species, and a cut-off of 95% for individual species. Pangenome collector’s curves were generated for each species group by calculating the number of unique genes present at increasing numbers of sampled genomes, with 1000 iterations of each sample size up to 250. Genetic clustering of genomes within species groups based on variable gene content was calculated and visualized using principal component analysis of accessory genes (PCA-A) using the R packages prcomp, vegan, and ggbiplot, with matrices of gene presence/absence used as input. Genes that were present in all isolates, present in only one isolate, or absent in only one isolate, were removed from analysis. PCA-A coordinate plots were visualized using GraphPad Prism version 7.0c.

### Core genome SNP comparisons, phylogenetic trees, evolutionary rate and recombination analyses

Within each genus, species, ST, or clade, SNPs were identified using snippy (https://github.com/tseemann/snippy). The most complete genome assembly (i.e. highest N50) was used as a reference genome for SNP analysis. Core genome SNPs, defined as SNPs at nucleotide positions shared across all genomes in the sample group being compared, were used to calculate pairwise SNP distances and to generate maximum likelihood phylogenetic trees. Trees were generated with RAxML v8.2 using the general time reversible model of evolution (GTRCAT), Lewis correction for ascertainment bias, and 100 bootstrap replicates^56^. Unless otherwise specified, reported SNP distances refer to core genome SNPs for all isolates belonging to the same ST. Pairwise SNP distances were visualized using the R package ggplot2. Recombination and evolutionary rates were calculated for STs in four species groups (*P. aeruginosa*, *Clostridioides difficile*, VRE and MRSA), and for STs within each group with more than 25 isolates. Estimates of relative recombination rates (R/Theta) and average size of recombinant sequences (delta) were assessed from core genome alignments using ClonalFrameML v1.12^24^ with default settings. The relative rate of recombination, which reflects the number of nucleotide changes introduced by recombination relative to each point mutation (r/m) was calculated as r/m = (R/Theta) × delta × ν^24^, where ν is the average distance between recombined sequences. A core genome alignment and recombination-corrected phylogenetic tree were used to estimate evolutionary rates using TreeTime^23^. Isolates that were found to be highly divergent from other isolates of the same ST (as revealed by an excess number of SNPs separating them from other isolates) were removed from the analysis.

### Antibiotic resistance gene detection and analysis

Acquired antimicrobial resistance genes were detected by querying genome assemblies against the ResFinder database using BLASTn^25^. A gene was considered present if the BLASTn percent identity multiplied by the sequence coverage was >80%. Resistance gene presence was mapped to a global phylogenetic tree constructed from amino acid sequences of 120 ubiquitous protein coding genes from the Genome Taxonomy Database Tool Kit^57^. Resistance gene co-occurrence was calculated using the %*% operator in R. This operator works by identifying the cross-products between any two genes found in a matrix of resistance genes identified in all isolates. The results were used to construct a relative frequency plot using the ggplot2 package in R. To include only the most frequently co-occurring gene pairs in the plot, a relative frequency of 80% and a combined frequency of 50% were used as cut-off thresholds. Additionally, genes found in >250 isolates were excluded as they were suspected of not being acquired resistance genes. ESBL and carbapenemase enzyme distributions were determined by assigning enzyme types based on protein sequence, and only 100% protein sequence matches are reported.

### Shared sequence detection and analysis

Putative mobile genetic elements were identified by searching for sequences >10kb that were present at high identity (>99.9%) in the genomes of isolates belonging to different species (<95% ANI) using nucmer^58^. Sequences were organized into clusters using all-by-all BLASTn v2.7.1^59^, and clusters were visualized with Cytoscape v3.8.2^60^. Clustered shared sequences were determined as resembling plasmids, insertion sequences (ISs), transposons, prophages, or integrative conjugative elements by BLAST against complete plasmids from NCBI databases^61^, MobileElementFinder^62^, PHASTER^63^, ProphET^64^ and ICEberg^65^, as well as comparison to the NCBI nr database and manual curation. Antimicrobial resistance genes in clustered sequences were identified by BLASTn against the ResFinder database^25^. Clusters of orthologous groups of proteins (COG) categories were assigned to genes present in one or more clustered sequences, and the distribution of genes in each COG category was visualized with the pie function in R.

## Data availability

Raw sequencing reads and genome assemblies were submitted to the NCBI Sequence Read Archive (SRA) and GenBank, with accession numbers listed in Table S1.

## Acknowledgements

We gratefully acknowledge all members of the Microbial Genomic Epidemiology Laboratory (MiGEL) at the University of Pittsburgh for their contributions to the EDS-HAT project. We also thank the Microbial Genome Sequencing Center (MiGS) for conducting Illumina sequencing of EDS-HAT isolates. Finally, we acknowledge Christi McElheny, Alina Iovleva, and Yohei Doi for assistance with ESBL typing. This work was supported by NIAID grants R21Al109459 and R01AI127472 to LHH, U01AI124302 to VSC, and by the Department of Medicine at the University of Pittsburgh School of Medicine (DVT). MMM was supported by grant KL2-TR001856. The funders had no role in study design, data collection and analysis, decision to publish, or preparation of the manuscript.

## Figure Legends

**Figure S1.**
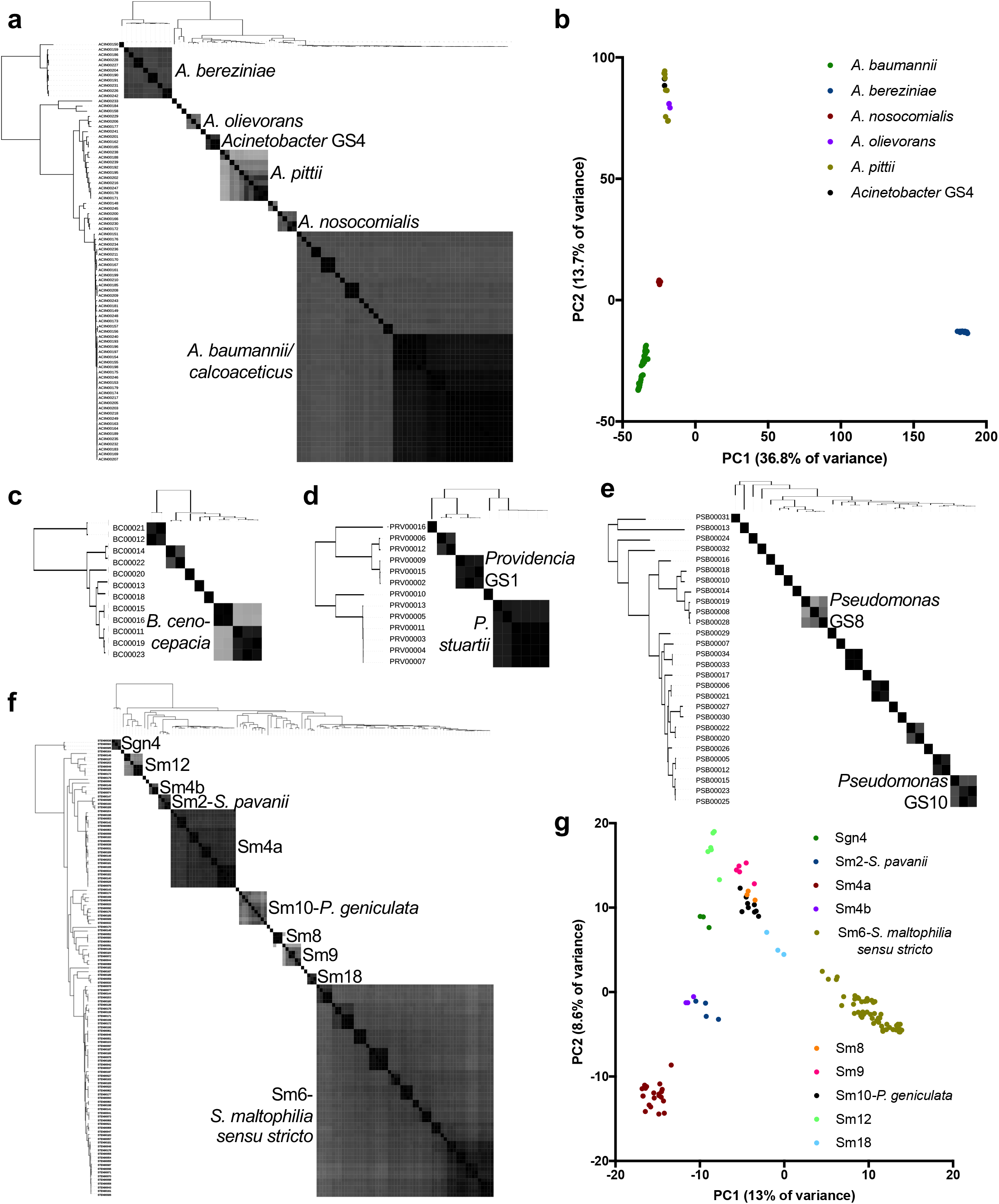
Average nucleotide identity (ANI) and principal components analysis of accessory genes (PCA-A) among diverse species groups sampled by EDS-HAT. (A) Phylogenetic tree with pairwise ANI values and (B) PCA-A plot for *Acinetobacter* spp. (C) Phylogeny and ANI of *Burkholderia* spp., (D) *Providencia* spp., (E) *Pseudomonas* spp., and (F) *Stenotrophomonas* spp. (G) PCA-A plot for *Stenotrophomonas* spp. Grey shading indicates ANI values >95%, with darker shading showing higher identity. PCA-A plots include species with >2 isolates.

**Figure S2.**
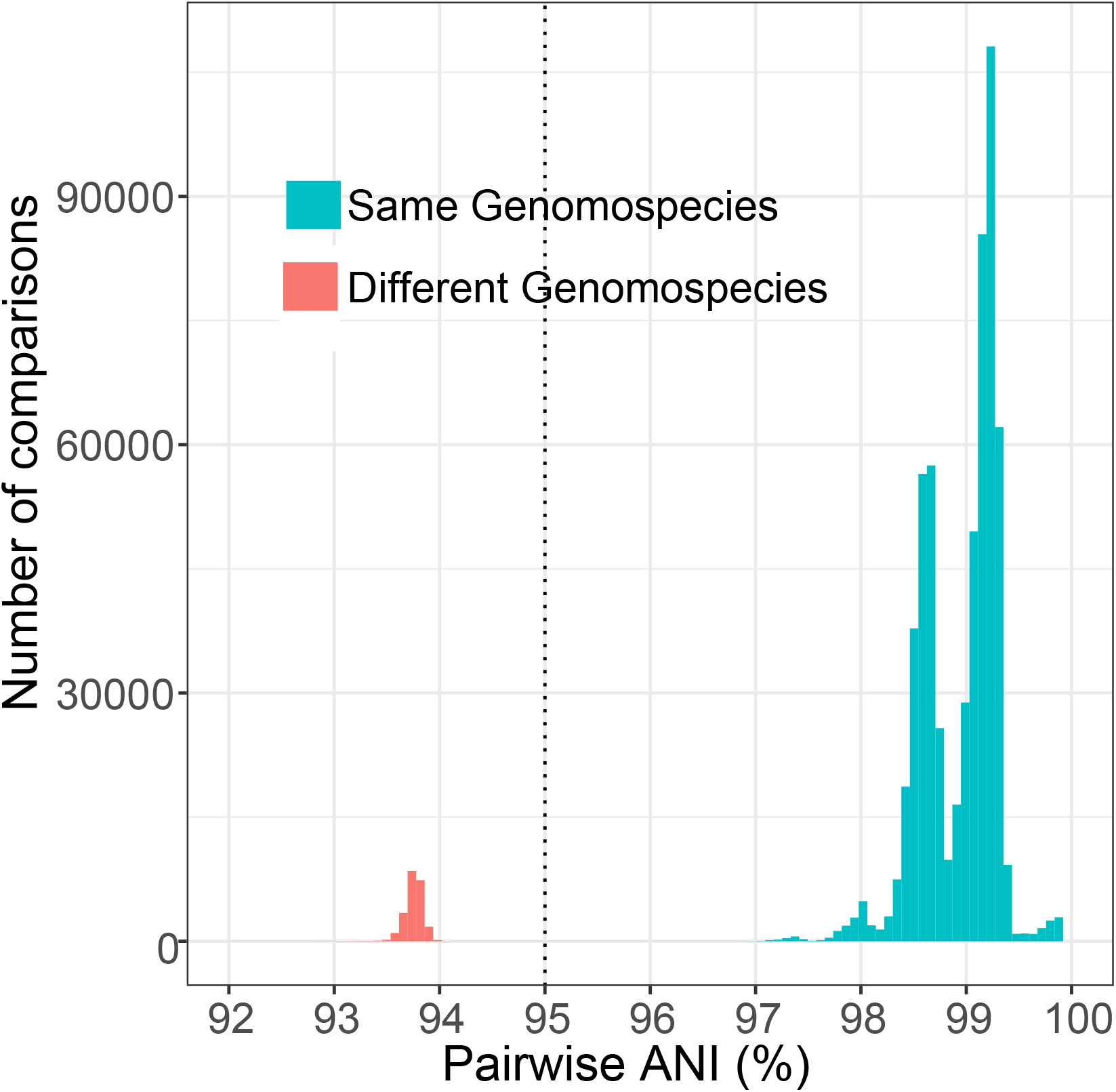
Average nucleotide identity (ANI) comparisons of *P. aeruginosa* isolates. Histogram of pairwise ANI values for 863 *P. aeruginosa* isolate genomes sampled by EDS-HAT. Dashed vertical line indicates 95% ANI. Comparisons in red are between isolates in *P. aeruginosa* Groups 1 or 2 versus isolates in the PA7-like Group 3, which appear to belong to a distinct genomospecies.

**Figure S3.**
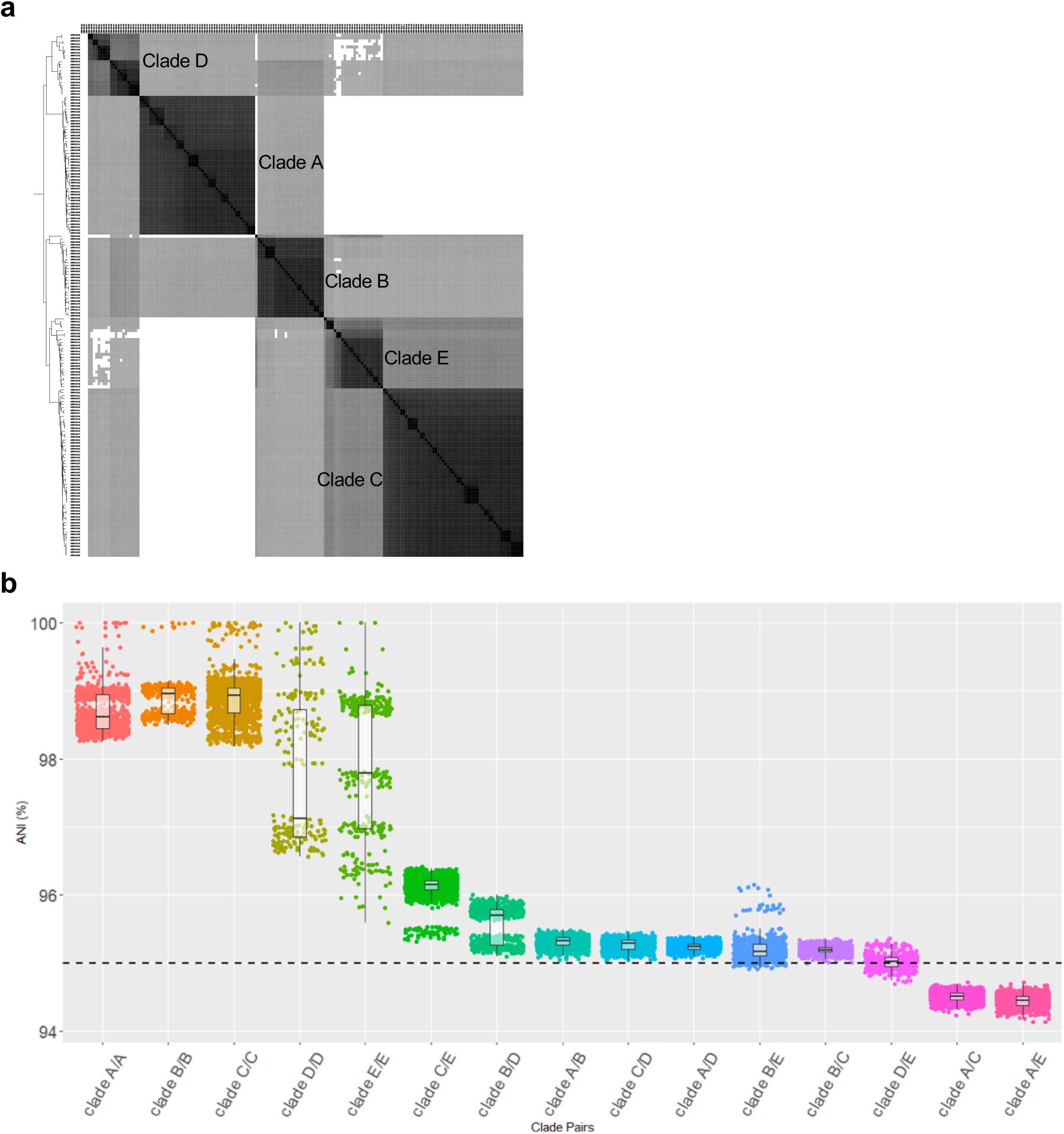
Average nucleotide identity (ANI) comparisons of *S. marcescens* isolates. (A) Phylogeny and ANI of 177 *S. marcescens* isolates sampled by EDS-HAT. Grey shading indicates ANI values >95%, with darker shading showing higher identity. White indicates ANI values <95%. (B) Distribution of pairwise ANI values for *S. marcescens* isolates belonging to the same or different clades, broken down into pairwise clade comparisons. All comparisons between isolates in Clade A vs. Clade C and Clade A vs. Clade E fall below the standard species cutoff of 95%.

**Figure S4.**
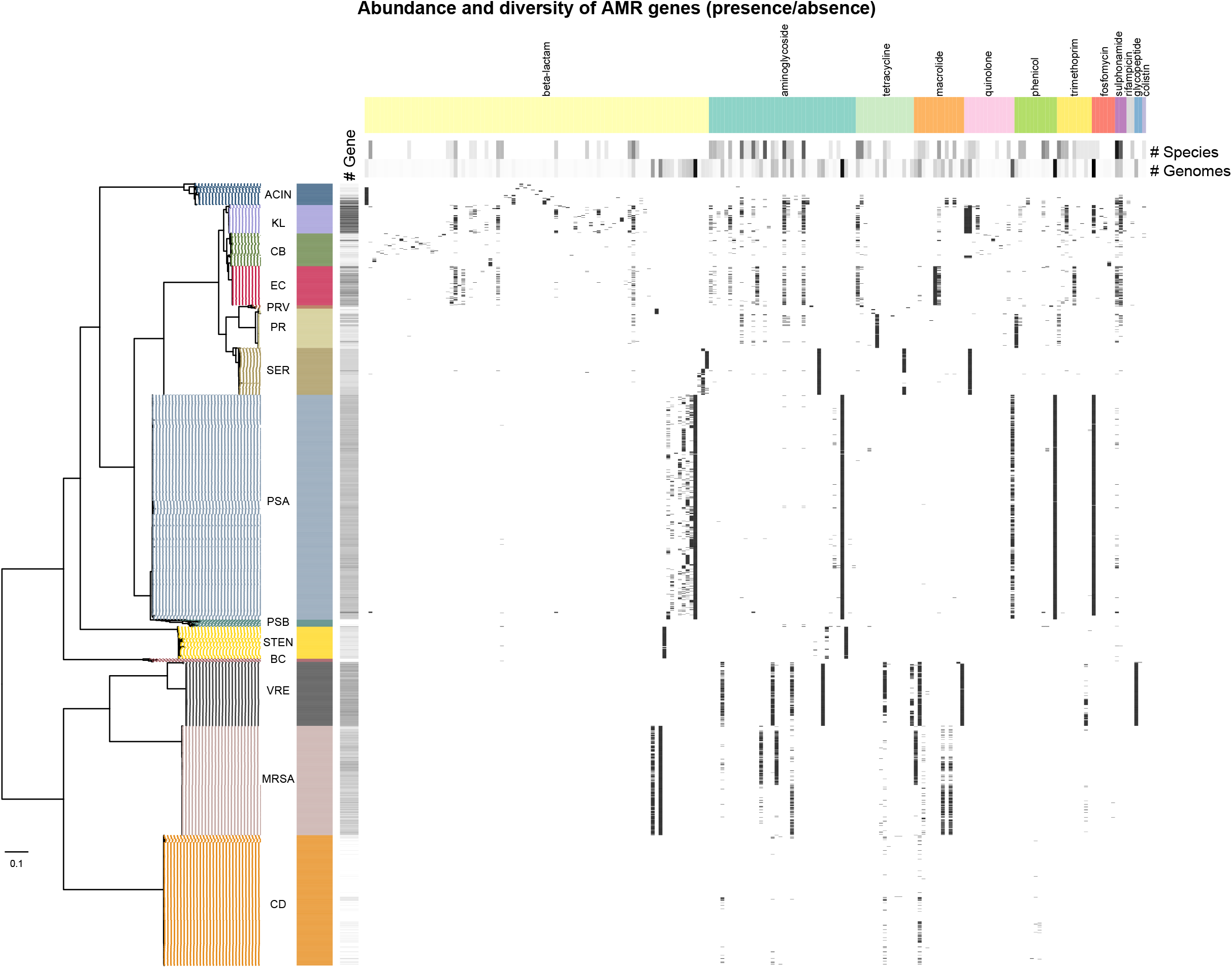
Distribution of antimicrobial resistance (AMR) genes among 3,004 clinical bacterial isolates from hospitalized patients. Resistance genes were identified by BLASTn comparison to the ResFinder database. Isolates are ordered according to their phylogenetic placement using the amino acid sequences of 120 ubiquitous protein-coding genes from the Genome Taxonomy Database Tool Kit. “# Gene” shows the number of AMR genes per genome, with darker shading indicating more AMR genes. The matrix shows the presence or absence of 202 AMR genes, grouped by antibiotic class. Heat maps at the top show the number of species groups and total number of genomes encoding each gene, with darker shading indicating higher numbers. Raw data used to make the matrix are available in Table S3.

